# An extinct clade sister to Eumetazoa: On the phylogeny of the Cambrian chancelloriids

**DOI:** 10.1101/2024.07.22.604532

**Authors:** Hao Yun, Xingliang Zhang, Glenn A. Brock, Jian Han, Luoyang Li, Bing Pan, Guoxiang Li, Joachim Reitner

## Abstract

The notable disparity of animal body plans can be traced back to the morphological innovations during the Cambrian explosion and represented by a number of exceptionally preserved soft-bodied and skeletal fossils that provide a compelling narrative for animal evolution. Chancelloriids, one of the extinct groups of Cambrian animals, have a distinctive body that characterized by a sclerite-bearing, flexible integument and a single apical opening leading into a central cavity devoid of unequivocal internal organs. Their phylogenetic position within the Metazoa, however, is controversial. Here, we describe new soft-bodied fossils of chancelloriids from the 518-million-year-old Chengjiang biota of China, which corroborate the unique bauplan pattern and reveal exquisite integument microstructures. The tiny protuberances and wrinkling structures of the integument are interpreted to be related to primitive epithelial contraction, suggesting that chancelloriids were a group of basal epitheliozoans and constitute an evolutionary clade that branched below all extant eumetazoans while above or close to the placozoans. Thus, the chancelloriid body plan likely filled one of the anatomical gaps between the Placozoa and the Eumetazoa.

---

The lineages of non-bilaterian basal metazoans, including Porifera, Cnidaria, Ctenophora and Placozoa, are remarkably disparate in anatomy and characterized by various asymmetrical or radially-symmetrical, diploblastic-grade (though questioned in poriferans) body plans (Martindale 2005; Dohrmann & Wörheide 2013; Dunn *et al*. 2015). In recent years, by means of a series of phylogenomic analyses, the relationship within Bilateria has been well deciphered, however, the placement of different basal metazoans that can largely reflect the early evolutionary history of animals are still in dispute (Martindale 2005; Dunn *et al*. 2008; Feuda *et al*. 2017; Pett *et al*. 2019). Hence, it is imperative to resort to fossil records (representing distinct lineages) of basal metazoans that reveal the historical dimension, in order to restore the picture for the early branching metazoan clades and fill the gaps of anatomical designs among extant taxa (Budd 2003).

The remarkable disparity of animal body plans can be traced back to the Ediacaran-Cambrian transition (560–510 Ma; namely Cambrian explosion) and represented by a number of exceptionally-preserved soft-bodied and skeletal fossils that provide a compelling narrative for the animal evolution (Shu 2008; Erwin *et al*. 2011; Erwin & Valentine 2013; Zhang & Shu 2021). Small shelly fossils (SSFs), dominated by non-bilaterian basal metazoans and protostomes, are particularly striking elements of the Cambrian Explosion; many of them once assigned to ‘Problematica’ or as bizarre disarticulated or cataphract elements from complex scleritomes, have led to heated debates regarding their biological affinities and phylogenetic positions (Bengtson & Missarzhevsky 1981; Bengtson *et al*. 1990; Qian *et al*. 1999; Bengtson 2005). Thus, integrated fossil evidence from isolated three-dimensional biomineralized sclerites along with superb articulated scleritome materials from the Konservat-Lagerstätten, is an effective way to accurately reconstruct the anatomy, morphology, and phylogenetic relationships of the enigmatic taxa (Conway Morris & Peel 1990; Bengtson 2005), such as the extinct Chancelloriida.

Chancelloriids were initially described as a family of the Porifera based on specimens from the Burgess Shale (Walcott 1920). Later, abundant and diverse isolated chancelloriid sclerites have been found in Cambrian SSFs worldwide (e.g., Sdzuy 1969; Luo *et al*. 1982; Bengtson *et al*. 1990; Qian *et al*. 1999; Yun *et al*. 2016; Yun & Zhang *et al*. 2019). Meanwhile, the family Chancelloriidae Walcott, 1920 was elevated to the order Chancelloriida Walcott, 1920 (Sdzuy 1969; Bengtson & Collins 2015). Although chancelloriids have long been interpreted as sponge or sponge-grade organisms due to their erected sac-like body, absence of organs, and sessile filter-feeding lifestyle (Walcott 1920; Sdzuy 1969; Rigby 1978; Cong *et al*. 2018), it is now clear that they had an epitheliozoan-grade flexible integument and a single, apical body opening (the apical orifice) (Bengtson & Hou 2001; Janussen *et al*. 2002; Randell *et al*. 2005; Bengtson & Collins 2015; Cong *et al*. 2018; Yun *et al*. 2018). The external calcareous (aragonitic) sclerites bedecked on the surface of integument possess relatively complex microstructures that resemble skeletal structures as seen in halkieriids, siphogonuchitids, and other ‘coeloscleritophorans’ (a group of ‘problematic’ bilaterians) (Bengtson & Missarzhevsky 1981; Bengtson 2005; Porter 2008). These similarities have complicated the phylogenetic identification of chancelloriids. In particular, the scleritome-bearing benthic vagrant animal halkieriids and their relatives have conclusively been recognized as stem or crown group lophotrochozoans (Conway Morris & Peel 1990; Conway Morris & Caron 2007) that possessed a body plan, trophic mode and life habit far removed from chancelloriids. Hence, chancelloriids were more plausibly proposed to be non-bilaterian basal metazoans (Bengtson & Collins 2015; Yun *et al*. 2018; Yun & Zhang *et al*. 2021), though their detailed zoological affinity is still in a state of flux.

Exceptionally preserved fossils of chancelloriids from the Cambrian Chengjiang biota (*ca*. 518 million years ago (Ma)) of China are described herein, which reveal distinctive epithelial microstructures and thus will help shedding new light on the phylogeny and early evolution of non-bilaterian basal metazoans.

## MATERIAL AND METHOD

### Fossil material

A total of 816 fossil specimens of two genera and four species (*Allonnia phrixothrix* Bengtson and Hou 2001, *Al. erjiensis* Yun et al. 2018, *Al. nuda* Cong et al. 2018, and *Dimidia simplex* Jiang in Luo et al. 1982) of chancelloriids were collected from five localities, including Sanjiezi (SJZ), Mafang (MF), Erjie (EJ), Jianshan (JS) and Shankou (SK) around the Kunming area, Yunnan Province, China (Table S1), all of them are deposited in the Shaanxi Key Laboratory of Early Life and Environments (LELE), Northwest University, Xi’an. The fossils are within the *Eoredlichia*-*Wutingaspis* Assemblage Biozone (*ca*. 518 Ma) of the Cambrian Yu’anshan Formation (Chengjiang biota) (Zhang *et al*. 2001; Yang *et al*. 2018).

### Fossil preparation and photography

Specimens were prepared and examined under a Nikon SMZ800 stereomicroscope with needles and brushes, when necessary; then photographed with a Canon EOS 5D Mark II camera under the magnesium light. Elemental distribution of specimens is analyzed by using Energy Disperse Spectroscopy (EDS) and X-Ray Fluorescence (XRF) under a FEI Q25 scanning microscope and a Bruker’s M4 TORNADO spectrometer, respectively.

### Cladistic analyses

Totally 26 taxa were involved in the analyses, in which Fungus is regarded as the outgroup and the rest 25 taxa include Choanoflagellata, Demospongiae, Hexactinellida, Homoscleromorpha, Calcarea, Placozoa, Anthozoa, Medusozoa, Priapulida, Onychophora, Phoronida, Brachiopoda, Arthropoda, Acoelomorpha, Annelida, Mollusca, Chordata, Echinodermata, *Halkieria* (representing the Cambrian Coeloscleritophora), and 6 chancelloriid genera *Chancelloria*, *Allonnia*, *Archiasterella*, *Nidelric*, *Dimidia*, and *Cambrothyra* (chancelloriids and *Halkieria* are fossil taxa). The analyses were performed in MrBayes 3.2.7 (Ronquist *et al*. 2012) and TNT 1.5 (Goloboff *et al*. 2008) by running a matrix with 117 characters (see Appendixes S1 and S2 for detail). The Bayesian phylogenetic analysis was performed using maximum parsimony algorithm with equal character weights and the maximum likelihood and gamma-distributed rate (mki + Γ) model. The TNT analysis was performed with a ‘traditional search’ algorithm.

## RESULTS

### Protuberances and wrinkling of the soft integument

Soft-bodied integuments of chancelloriids are preserved as compressed films of clayey minerals and iron oxides highlighted by a reddish colour (Figs. 1, S1, S2). There are distinctive tiny protuberances ornamented on the surface of the integument, especially in *Al. phrixothrix* (Fig. 1C–E, G). The size of protuberances is consistent throughout the individual body but varies across different taxa. For instances, protuberances are 40–50 μm in diameter in the integument of *Al. erjiensis* (120–150 mm in height) (Yun *et al*. 2018), while they are 120–180 μm in diameter in *Al. phrixothrix* (150–300 mm in height) (Fig. 1; see also Bengtson & Hou 2001; Janussen *et al*. 2002), 60–80 μm in *Archiasterella coriacea* (about 110 mm in height) (Bengtson & Collins 2015), and about 80 μm in *Ar. fletchergryllus* (70–50 mm in height) (Randell *et al*. 2005).

**FIG. 1.**
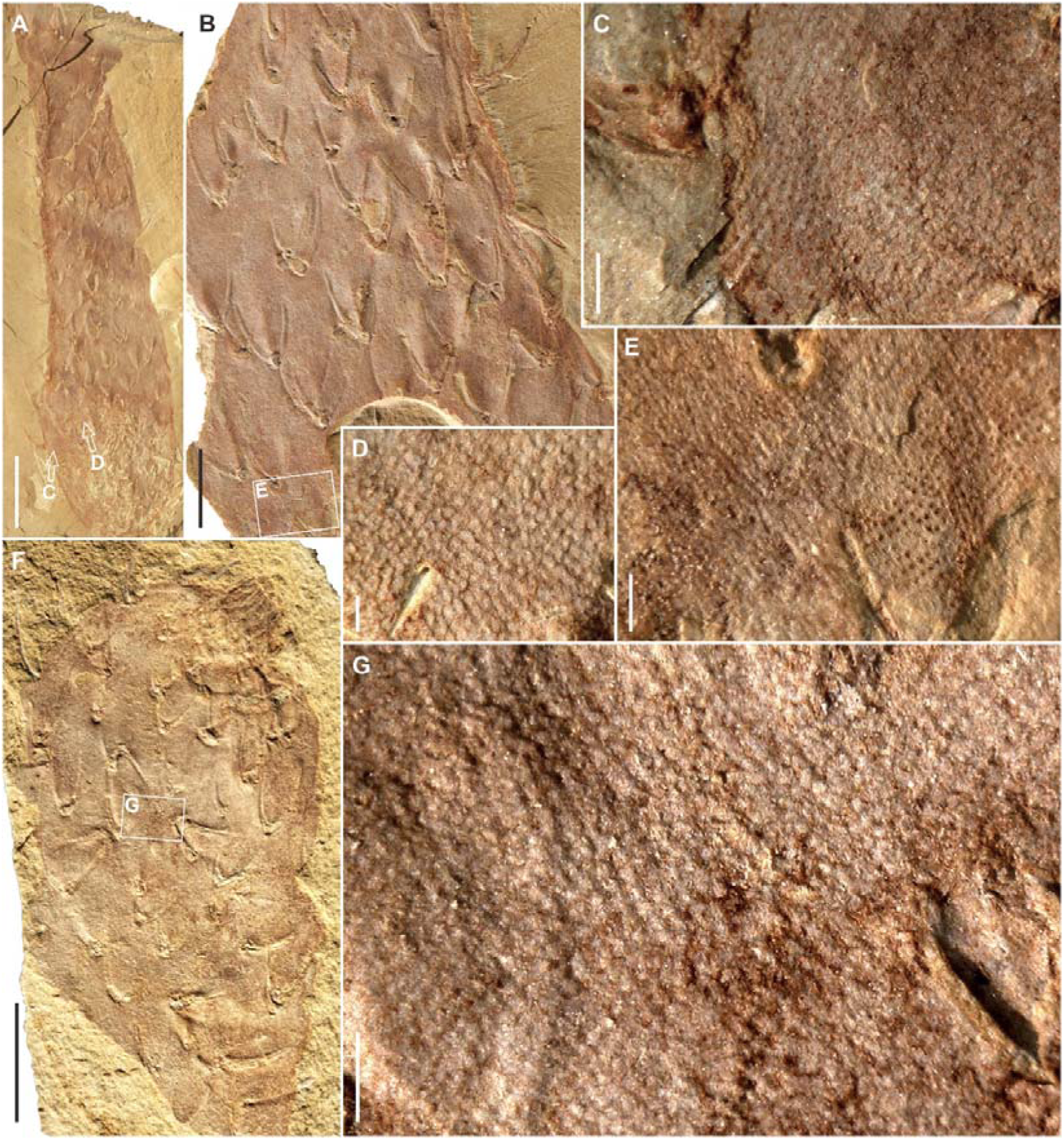
Microstructures of the chancelloriid integument. A, B, *Allonnia phrixothrix* from the Chengjiang biota, specimens SJZ-B03-179A and SJZ-B03-179B, respectively, showing highlighted reddish colour soft integument. C, D, details of A, showing remarkable protuberances. E, detail of B, showing protuberances and wrinkles (protuberances distributed in distinct rows, forming a series of parallel stripes). F, *Al. phrixothrix* from the Chengjiang biota, specimen SJZ-B03-180A. G, detail of F, showing distinctive protuberance. Scale bars represent: 10 mm (A, F), 5 mm (B), 2 mm (C, E, G), 1 mm (D).

**FIG. 2.**
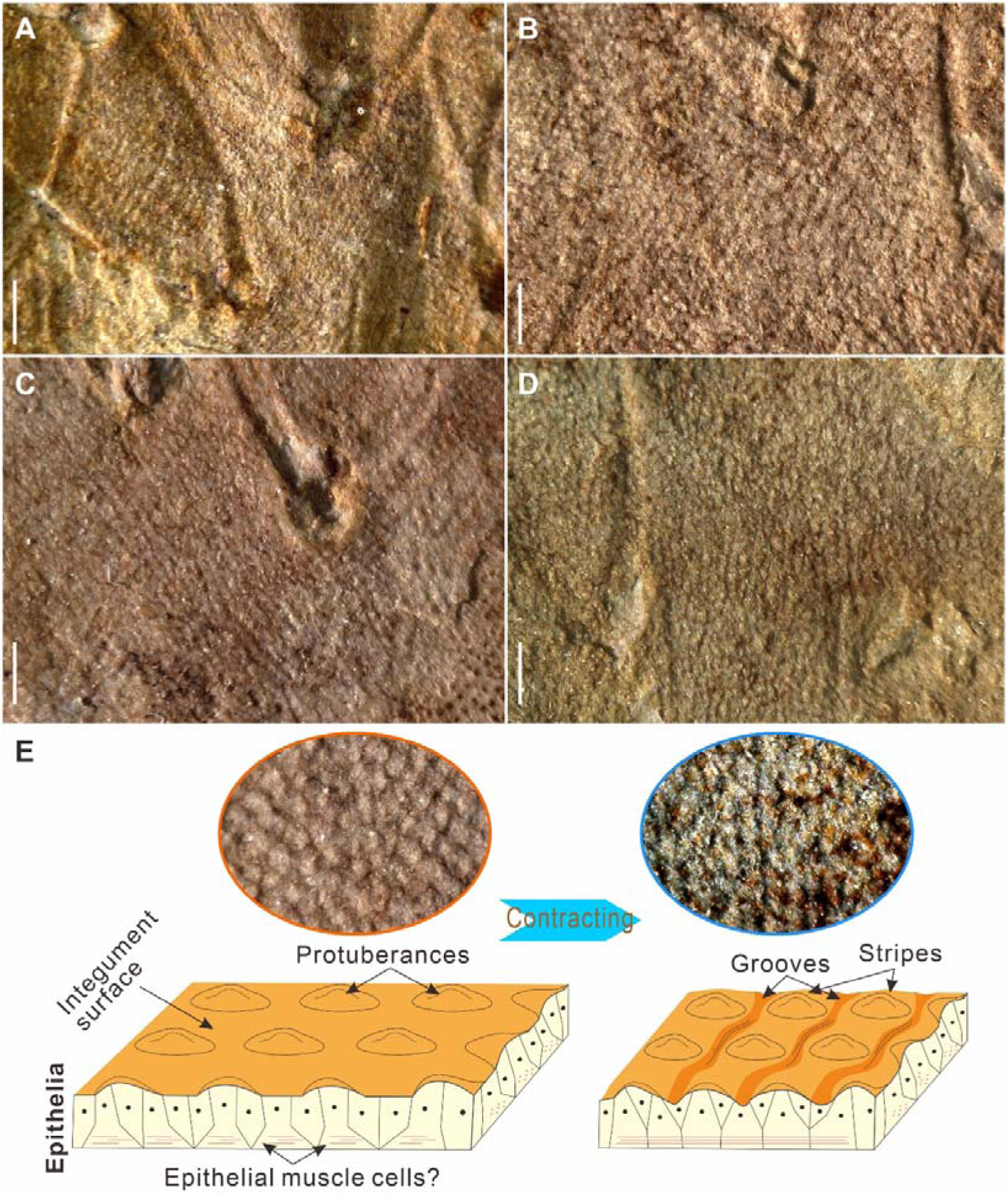
Wrinkling on the chancelloriid integument. A–D, integuments of the Allonnia phrixothrix from the Chengjiang biota, showing wrinkling microstructures; A, detail of the specimen SJZ-B04-095B (Fig. S1A); B, detail of the specimen SJZ-B03-180A (Fig. 1F); C, detail of the specimen SJZ-B03-179B (Fig. 1B); D, detail of the specimen SJZ-B05-010A (Fig. S2A). E, interpretation of the wrinkling microstructures formed by couplets of parallel stripes (rows of protuberances) and grooves. Scale bars represent: 2 mm.

The protuberances in the fossil specimens studied herein also reveal that these microstructures can be regularly arranged in rows, forming parallel, stripe-like structures that mostly distributed in the middle and lower part of the body (Figs. 1E, 2, S2). The rows or stripes are subparallel or diagonal to the vertical axis of the body (for example in *Al. phrixothrix*) and each is about 150–180 μm in width (generally accordant to the diameter of protuberances) and spaced by parallel grooves that are 80–100 μm wide (Fig. 2A–D). The compact couplets of stripes and grooves are morphologically reminiscent of wrinkling on the soft integument.

### Phylogenetic analyses

A detailed survey on chancelloriid fossils including both complete bodies and isolated sclerites provides a dataset (Data S1) for facilitating morphological comparisons and tentative phylogenetic (cladistic) analyses on different chancelloriid genera and their general body plan within a broad spectrum of animal groups. The main analysis is based on 117 characters across 25 animal taxa with Fungi as the outgroup (Appendix S1). The Ctenophora is excluded herein in consideration of the hot debates on its molecular and possible ‘hidden’ biology that hardly reflected on the anatomy of fossils (e.g., Dunn *et al*. 2008; Philippe *et al*. 2009; Dunn *et al*. 2015; Feuda *et al*. 2017; Laumer *et al*. 2019; Pett *et al*. 2019; Burkhardt *et al*. 2023). A consensus tree was resolved from credible sets of 659 sampled trees in the Bayesian analysis and 3 maximum parsimonious trees (tree length = 148; total fit = 93.8; adjusted homoplasy = 7.2, for all the trees) and a strict consensus tree of them were resolved in the TNT analysis. These trees indicate that 1) the Chancelloriida, represented by *Chancelloria*, *Archiasterella*, *Allonnia*, *Dimidia*, *Nidelric*, and *Cambrothyra*, forms a monophyletic clade sister to (or as a stem group of) the Eumetazoa comprising Cnidaria and Bilateria, and 2) within the chancelloriid group, *Cambrothyra* was diverged before the rest genera (Figs. 3, S3, S4).

**FIG. 3.**
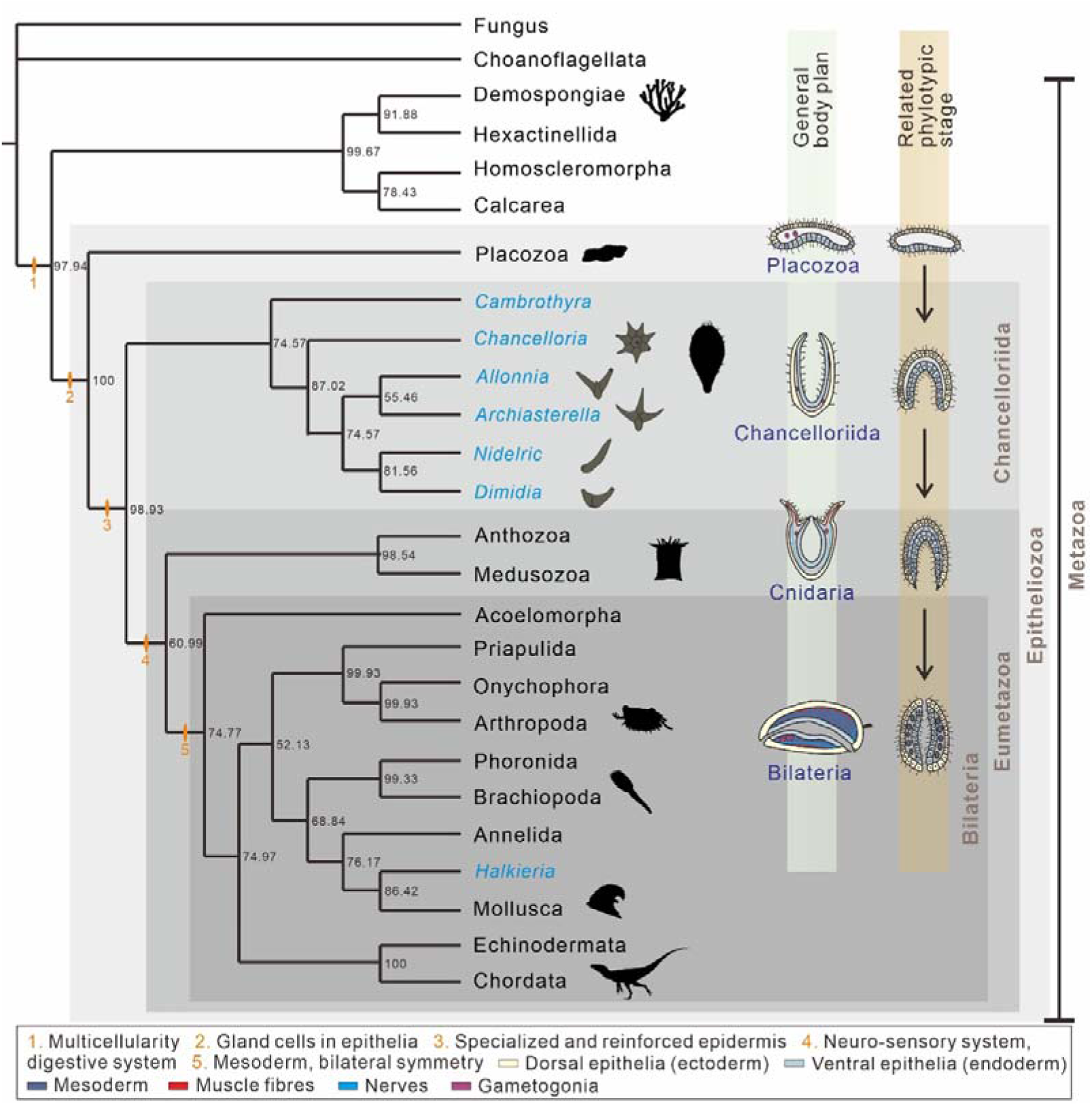
Morphological-based cladistic analysis and an evolutionary scenario of the epitheliozoan body plan. The consensus tree is obtained by using Bayesian phylogenetic analysis based on a dataset containing 26 taxa and 117 characters. Numbers at the nodes are percentages of posterior probabilities. The orange marks and numbers in the tree indicate synapomorphies as well as changes affecting main body plan characters of specific clades. Silhouettes of the representative animal clades are from *PhyloPic* (www.phylopic.org), except the one for chancelloriids. Within the clade of chancelloriids, the grey sketches are typical sclerite forms of each genus (Fig. 4). The gross evolutionary scenario of the animal body plan is loosely based on the developmental stages of metazoan embryogenesis (Martindale 2005; Schierwater *et al*. 2009), showing a continuous sequence of the body plan complexity from Placozoa to Bilateria.

**FIG. 4.**
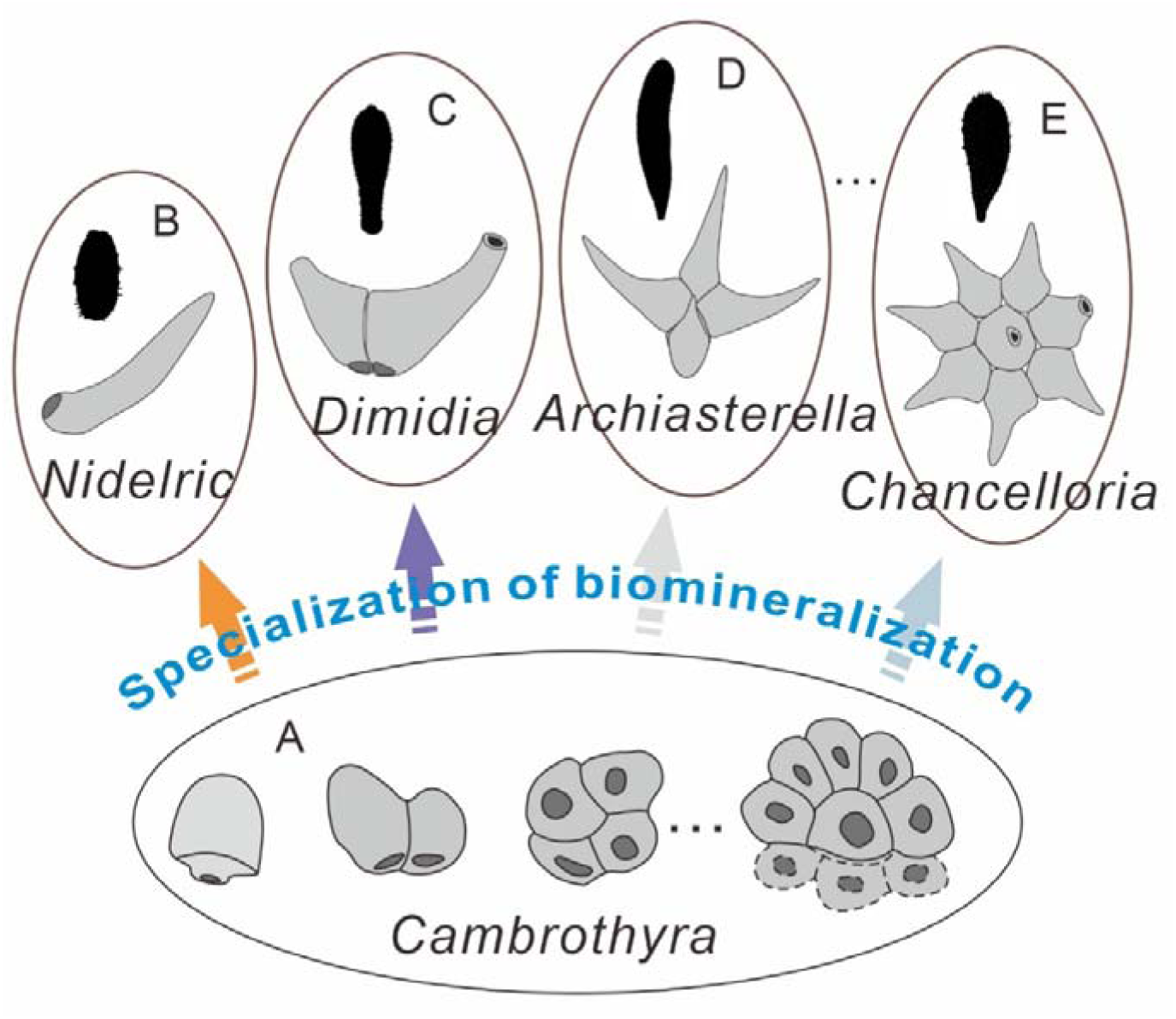
Specialization of biomineralization in chancelloriids. A, different forms of the *Cambrothyra* sclerites. B–E, typical sclerite structures and body shape silhouettes of the chancelloriid genera *Nidelric*, *Dimidia*, *Archiasterella*, and *Chancelloria*, respectively.

Furthermore, an additional analysis was performed based on a slightly modified character matrix, a ‘conservative’ version with more “?” coding for chancelloriids (see Appendix S2 for detail), from the one in the main cladistic analysis. The analysis operated in TNT 1.5 (with a ‘traditional search’ algorithm) resolves three maximum parsimonious trees (tree length = 148; total fit = 93.8; adjusted homoplasy = 7.2) and a strict consensus tree of them (Fig. S5). The topologies of the maximum parsimonious trees are the same to the counterparts resolving in the main analysis, while in the strict consensus tree, Chancelloriida, Cnidaria, and Bilateria are clades from a tripartite bifurcation that sister to Placozoa. These results also support that chancelloriids are basal epitheliozoans and probably a stem-group of the Eumetazoa.

## DISCUSSION

### Skeletal structure and chancelloriid ancestry

Besides supporting the body, a part of the integument (epidermis) in chancelloriids was an organic template that served in sclerite biomineralization (Bengtson & Hou 2001; Yun & Zhang *et al*. 2021). Recent discoveries reveal that the biomineralization of the external skeleton of chancelloriids involved a close cooperation between numerous cell types facilitating a suite of complex processes, such as secreting, integrating, and distributing fibrous aragonitic biominerals, to create composite, hollow sclerites (Mehl 1996; Bengtson & Hou 2001; Porter 2008; Yun & Cui *et al*. 2021; Yun & Zhang *et al*. 2021). Therefore, cells (and tissues) participated in these processes, that together with the organic template, formed a specialized epithelio-biomineralizing structural unit, which is functionally similar to related structures found in a range of eumetazoans, such as Cnidaria, Mollusca, and Hyolitha (Bengtson 2005; Schmidt-Rhaesa 2007; Erwin & Valentine 2013; Yun & Zhang *et al*. 2021).

As products of the epithelio-biomineralizing structural unit, the sclerites of chancelloriids are constructed by different numbers of articulated hollow rays (Qian & Bengtson 1989; Yun & Zhang *et al*. 2019). In most chancelloriid genera, such as *Chancelloria*, *Archiasterella*, *Allonnia*, and *Dimidia*, sclerites are composite in structure with at least two conical rays. (Janussen *et al*. 2002; Randell *et al*. 2005; Bengtson & Collins 2015; Yun *et al*. 2018; Yun & Zhang *et al*. 2019; Peng *et al*. 2024; Yun *et al*. 2024). However, in the genus *Nidelric*, only revealed in the Chengjiang and Guanshan biotas (Cambrian Stages 3 to 4; *ca*. 518–510 Ma) of China, single-rayed sclerites (not composite forms) are dominated the scleritome (Hou *et al*. 2014; Zhao *et al*. 2018). In consideration of comprehensive fossil records, the oldest known chancelloriids date back to the beginning of Cambrian (*ca*. 535 Ma) and belong to *Chancelloria* as well as a specialized group named *Cambrothyra* (Qian & Zhang 1983; Steiner *et al*. 2004; Moore *et al*. 2010). The basal elements of *Cambrothyra* sclerites are hollow and broadly ellipsoid in shape since their conical rays are very small and reminiscent of buds (Fig. 4A). There are a series of combination forms of these ellipsoid elements (all from the same sampled horizon), including single, double, quadruple, and special types composed of six to nine lateral elements surrounding one central element (Qian & Zhang 1983; Moore *et al*. 2010), which closely correspond to the characteristic sclerite forms of *Nidelric*, *Dimidia*, *Archiasterella*, and *Chancelloria*, respectively, in general structure (Fig. 4). Consequently, the putative complete scleritome of *Cambrothyra* resembles a biomineralizing ‘prototype’ that comprises most kinds of sclerite structures subsequently developed in the scleritomes of specific chancelloriid genera.

It is widely accepted that animals with biomineralized skeletons have rapidly (within 40 Ma during the Ediacaran-Cambrian transition) evolved from unmineralized ancestors by co-option of molecules (gene suites and proteins) that earlier served other biological functions (Wood & Zhuravlev 2012; Murdock 2020; Zhang & Shu 2021). Following this evolutionary trajectory, the biomineralizers trended toward an increasing control of secreting tissues and biominerals (Kocot *et al*. 2016; Murdock 2020; Gilbert *et al*. 2022). Although the unmineralized ancestor of chancelloriids is still to be discovered, the general form and relatively simple shape of sclerites in *Cambrothyra* likely represent a nascent stage of biomineralization when chancelloriids had less control on secretory skeletal systems (Moore *et al*. 2014). The scleritomes of other chancelloriid genera, each dominated by a specific form of sclerites, indicate increasing proficiency and specialization on constructing the biomineralized framework. Thus, *Cambrothyra* is likely to represent a key ancestor in the evolution of all chancelloriids.

### Body plan and phylogenetic position

The tubercular microstructures on the integument surface of chancelloriids described above have also been reported as small, plate-like ornamentations (Bengtson & Hou 2001; Randell *et al*. 2005; Bengtson & Collins 2015) or reflections of underlying epithelial cells (Janussen *et al*. 2002) that distributed across the genera *Allonnia*, *Archiasterella*, and *Chancelloria.* The three-dimensional preservation and occasional imbricated arrangement of the protuberances contradict the cell impression interpretation (Randell *et al*. 2005; Bengtson & Collins 2015) but emphasize the strengthening of the epidermis. The couplets of stripes and grooves (formed by protuberances) on the integument are firstly revealed herein, which are regularly distributed (mostly in the middle and lower part of the body surface) and orientated (diagonal or parallel to the vertical body axis), reflecting an original wrinkling structure rather than a taphonomic artefact. Therefore, the chancelloriid epidermis is thick enough to be corrugated and likely contracted when compare the wrinkling to the homogenously distributed protuberances (Fig. 2E). Namely, the chancelloriid body could contract in response to external physical or biological stimuli, so that adjacent rows of protuberances draw close to form couplets of stripes and grooves. Such process is likely related to primary epitheliomuscular cells (EMCs) (Schmidt-Rhaesa 2007) or contractile myoepithelial cells that act in the context of epithelial contraction (Brunet & Arendt 2016).

It is well accepted that chancelloriids have a subcylindrical body consisting of a flexible integument bedecked by biomineralized sclerites with variable numbers of rays, and a single apical opening (orifice) leading into a central cavity without any unequivocal internal organs (Bengtson & Hou 2001; Bengtson & Collins 2015; Cong *et al*. 2018; Yun *et al*. 2018). The body usually tapers both apically and abapically, forming a constricted apex with an obconical basal end anchored to the seafloor substrate, suggesting that this animal was sessile and upright in life position (Kloss *et al*. 2009; Bengtson & Collins 2015). The apical orifice is surrounded by a palisade-like tuft formed by a suite of modified single-rayed sclerites (Figs. 1F, 5, S1) (Bengtson & Collins 2015; Yun *et al*. 2018; Yun & Brock *et al*. 2019), which likely functioned in obstructing large particles or predators from invading the body cavity. These modified sclerites are basically parallel to each other, with their inflated proximal bases fused or articulated together to provide a rigid tuft structure (Yun & Brock *et al*. 2019).

**FIG. 5.**
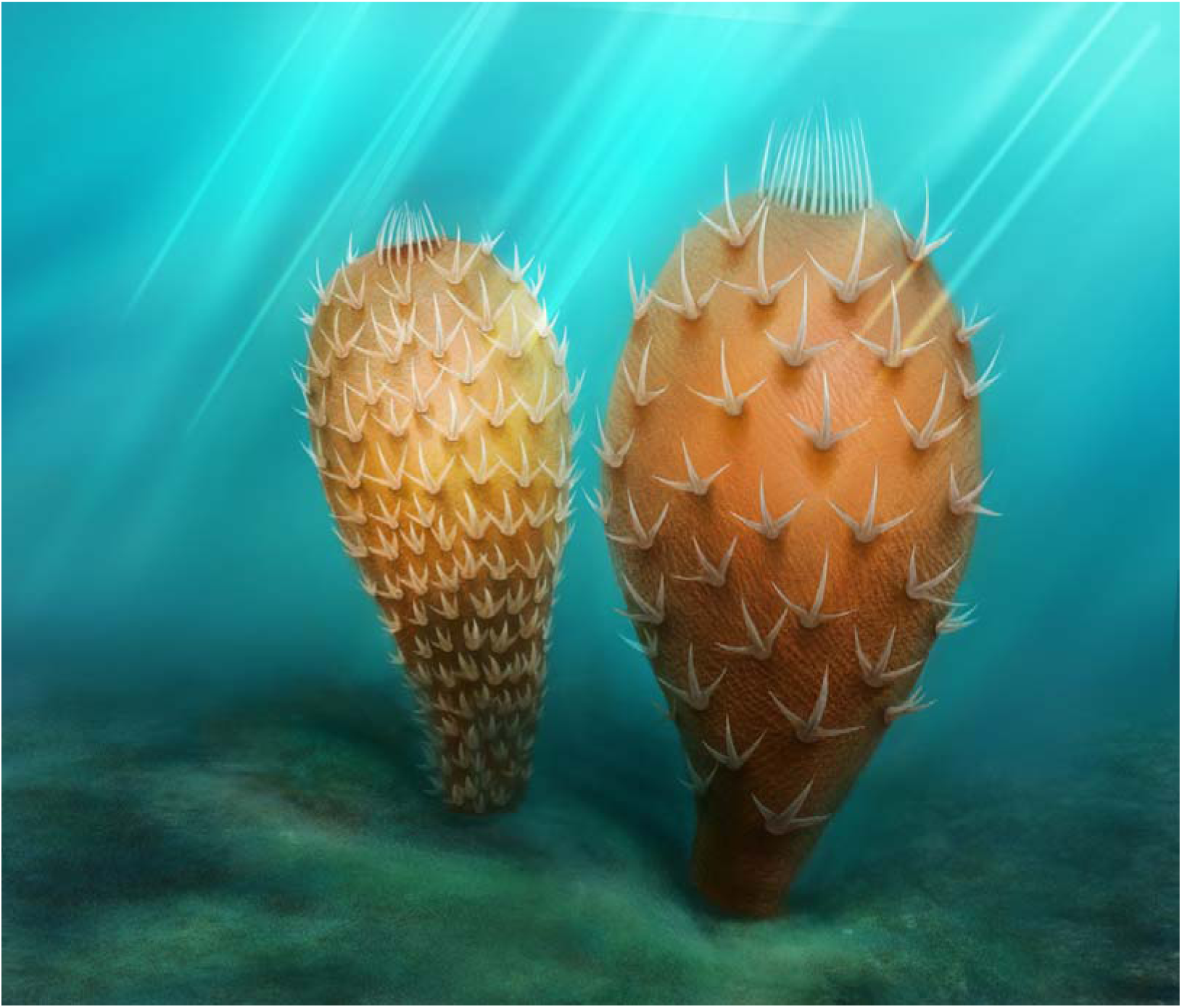
Reconstruction of representative chancelloriids from the Chengjiang biota. Artworks of *Allonnia erjiensis* (left) and *Al. phrixothrix* (right) are drawn by Xi Liu; copyright Northwest University.

Apart from a hypothesis that chancelloriids may acquire nutrition from bacterial or algal symbionts in the internal cavity (Bengtson & Hou 2001; Bengtson & Collins 2015), a sedentary filter feeding strategy is more reasonable due to the lack of tentacles or any other feeding organs. However, there is no evidence for a water flow through system that characterize Porifera, because of the absence of ostia-like openings in the body wall. Therefore, the apical orifice, as the only opening in the chancelloriid bauplan, potentially functioned as both inhalent and exhalent openings for water exchange between the inner cavity and environment (Bengtson & Collins 2015; Yun *et al*. 2018; Zhao *et al*. 2018). This suggests that chancelloriids employed body expansion and contraction relaying on at least the existence of abovementioned EMCs (Bengtson & Collins 2015; Yun *et al*. 2018).

In summary, the body plan of chancelloriids reveals a peculiar anatomy and form disparate from any extant metazoan phylum:

1. Chancelloriids have a sedentary, sac-like body reminiscent of sponges, but the flexible, corrugated body wall (epidermal system) with distinctive surface protuberances and microstructures and a sophisticated biomineralizing structural unit is radically different from and more complicated than the organization of bauplan in Porifera. Although extant homoscleromorph sponges possess a basement membrane, their ‘epidermis’ still refers to the thin and fragile choanoderm and pinacoderm (Gazave *et al*. 2012).
2. Chancelloriids have an epidermal organization congruent with placozoans and eumetazoans. Further, the relatively large and erected body shape, biomineralizing structural unit with differentiated cell types and tissues, and probable EMCs form a bauplan more complicated than Placozoa, since all placozoans (three species described to date) have a minute flat body that composed only of six types of cells (Srivastava *et al*. 2008; Smith *et al*. 2014; Schierwater *et al*. 2021a, 2021b).
3. Chancelloriids are characterized by a series of hollow aragonitic sclerites with regular fibrous microstructures that also occurs in many skeletal eumetazoans, such as Cnidaria and Mollusca (Carter 1990; Qian *et al*. 1999; Li *et al*. 2019). However, it is noted that this type of biomineralizing pattern could have been acquired independently by co-option of a similar ‘biomineralization toolkit genes’ across different lineages during the Cambrian explosion (Murdock 2020; Yun & Zhang *et al*. 2021; Gilbert *et al*. 2022). The absence of feeding tentacles, internal organs, or a certain nerve system distances chancelloriids from definite Eumetazoa.

### Implications for the evolution of epitheliozoans during the Cambrian explosion

Epitheliozoa is an evolutionary clade consisting of Placozoa and Eumetazoa (Ax 1996; Dohrmann & Wörheide 2013). As mentioned above, Placozoa has a minute flat body composed of six types of cell (ventral epithelial cell, dorsal epithelial cell, gland cell, lipophil cell, fiber cell, and crystal cell) (Smith *et al*. 2014). Comparatively, Eumetazoa is characterized by further evolving muscle cells and a neuro-sensory system (Martindale 2005; Philippe *et al*. 2009). Although it is not possible to compare cell-to-cell level traits, chancelloriids, possessing no obvious diverticular features in the central cavity or a preserved nervous system, but developed an epidermis (and probable EMCs) and a relatively complex process of sclerite biomineralization that fits within the concept of a basal epitheliozoan that branched between Placozoa and crown-group Eumetazoa.

Chancelloriids are morphologically more complex than placozoans mainly due to their erected body shape and well-developed aragonitic skeletons produced by a specialized biomineralizing structural unit. While it is noted that despite the apparent cellular and organismal simplicity, the genome of placozoans shows a higher diversity than that of sponges and encodes a rich array of transcription factors and signaling pathways that are associated with eumetazoan developmental processes (Schierwater *et al*. 2008; Srivastava *et al*. 2008; Schierwater *et al*. 2021a, 2021b). The epidermis of placozoans can be subdivided into two cellular layers; the lower layer contains ciliated ventral epithelial cells, gland cells, and lipophil cells that function in digesting food (mainly microorganisms), and the upper layer is composed of dorsal epithelial cells (Smith & Mayorova 2019). In between the epithelia are fibre cells and crystal cells; the latter are arrayed around the perimeter of the animal and each of them contains an aragonitic crystal that possibly related to sensing gravity (Mayorova *et al*. 2018). Therefore, it is not unreasonable to deduce a chancelloriid-like body plan derived from an ancestral placozoan, but with the invagination (and differentiation) of epidermal layers and a reversal in the direction of growth. The development of biomineralizing capability may originates from the aragonite-favoured habit (of the crystal cells) as well as the co-option of ‘biomineralization toolkit genes’ to encode related regulatory proteins, such as ACC (amorphous calcium carbonate) binding proteins and calcium metabolism-associated proteins (Zhang *et al*. 2019), during the Ediacaran-Cambrian transition. Furthermore, the potential EMCs in chancelloriids indicates that these animals probably formed a distinctive early-branched clade of stem group of Eumetazoa, which represents an extinct ‘intermediate’ type within an early evolutionary trajectory of epitheliozoans that loosely based on the embryogenesis (phylotypic stages) of different metazoan clades (Martindale 2005; Schierwater *et al*. 2009; Švorcová 2012; Galis & Metz 2021) (Fig. 3).

The Cambrian explosion records a rapid change in morphospace and the evolutionary landscape associated with an episodic increase in biomineralization across phyla resulting in and an increasing of ecosystem complexity (Erwin & Valentine 2013; Zhang & Shu 2021; McMenamin 2023). And most interestingly, the Cambrian animal world is characterized by co-occurrence of abundant stem-groups and crown-groups within different lineages of metazoans (Budd 2003; Erwin *et al*. 2011; Erwin & Valentine 2013). There were a number of class- and phylum-levelled stem-group taxa with regarding to the Linnaean classification, such as Archaeocyatha (an extinct class of Porifera or an ‘intermediate’ metazoan clade between Porifera and Eumetazoa) (Hooper & van Soest 2002; McMenamin 2023; Wang *et al*. 2024), Hyolitha (an extinct class of Mollusca or a phylum of lophotrochozoans) (Malinky & Yochelson 2007; Liu *et al*. 2020), and Vetulicolia (an extinct phylum of deuterostomes) (Shu *et al*. 2001). Chancelloriids described herein, along with these intriguing Cambrian body plans, represent extinct morphological innovations of early metazoans (Gould 1989; Budd 2003; Erwin *et al*. 2011; Erwin & Valentine 2013). On a hindsight view, although chancelloriids and most of these stem-group taxa totally went extinct not beyond the Cambrian Period (except for hyoliths, which existed within the Palaeozoic) (Shu 2008; Erwin & Valentine 2013; Zhang & Shu 2021), their explorations on the body plan of epitheliozoans have filled significant morphological gaps among early branches of the evolutionary tree of animals.

## Acknowledgements

We are grateful for the fieldwork and technical assistances from Cong Liu, Jie Sun, Meirong Cheng, and Juanping Zhai. The reconstruction artwork was drawing by Xi Liu. Thanks go to reviewers and editors for constructive comments.

## Author contributions

**Conceptualization** HY, XZ; **Data curation** HY; **Formal Analysis** HY; **Funding Acquisition** HY, XZ; **Investigation** HY, XZ, GAB, JH, LL, BP; **Methodology** HY, JH; **Project Administration** HY, XZ; **Resources** HY, XZ; **Supervision** XZ, GAB, JR; **Validation** HY, XZ, GL; **Visualization** HY; **Writing – Original Draft Preparation** HY, XZ, LL; Writing – Review & Editing HY, XZ, GAB, JH, LL, BP, GL, JR.

