## Appendix; Data for "An extinct clade sister to Eumetazoa: On the phylogeny of the Cambrian chancelloriids": Supp.file.1.docx

**This file includes:**

Table S1

Figures S1 to S5

**Other supplementary files (separate files) include:**

Appendix S1. Description on the character matrix for the cladistic analyses.

Appendix S2. Character modifications in the matrix for the additional cladistic analysis

Data S1. Records of current known chancelloriid fossils.

Data S2. Character matrix for the main cladistic analysis.

Data S3. Character matrix for the additional cladistic analysis.

**TABLE S1.** Quantity statistics of studied chancelloriids from the Cambrian Chengjiang biota in Kunming area, Yunnan Province, China.

| **Locality** | ***Allonnia*** | | | ***Dimidia simplex*** | **Sum** |
| --- | --- | --- | --- | --- | --- |
|  | ***Al*. *phrixothrix*** | ***Al*. *erjiensis*** | ***Al*. *nuda*** |  |  |
| Sanjiezi (SJZ) | 425 | 34 | 17 | 14 | 488 |
| Mafang (MF) | 78 | 10 | 13 | 2 | 103 |
| Jianshan (JS) | 88 | 21 | 14 | 3 | 126 |
| Erjie (EJ) | 58 | 1 | 4 | 1 | 64 |
| Shankou (SK) | 25 | 7 | 0 | 3 | 35 |
| Sum | 672 | 73 | 48 | 23 | 816 |

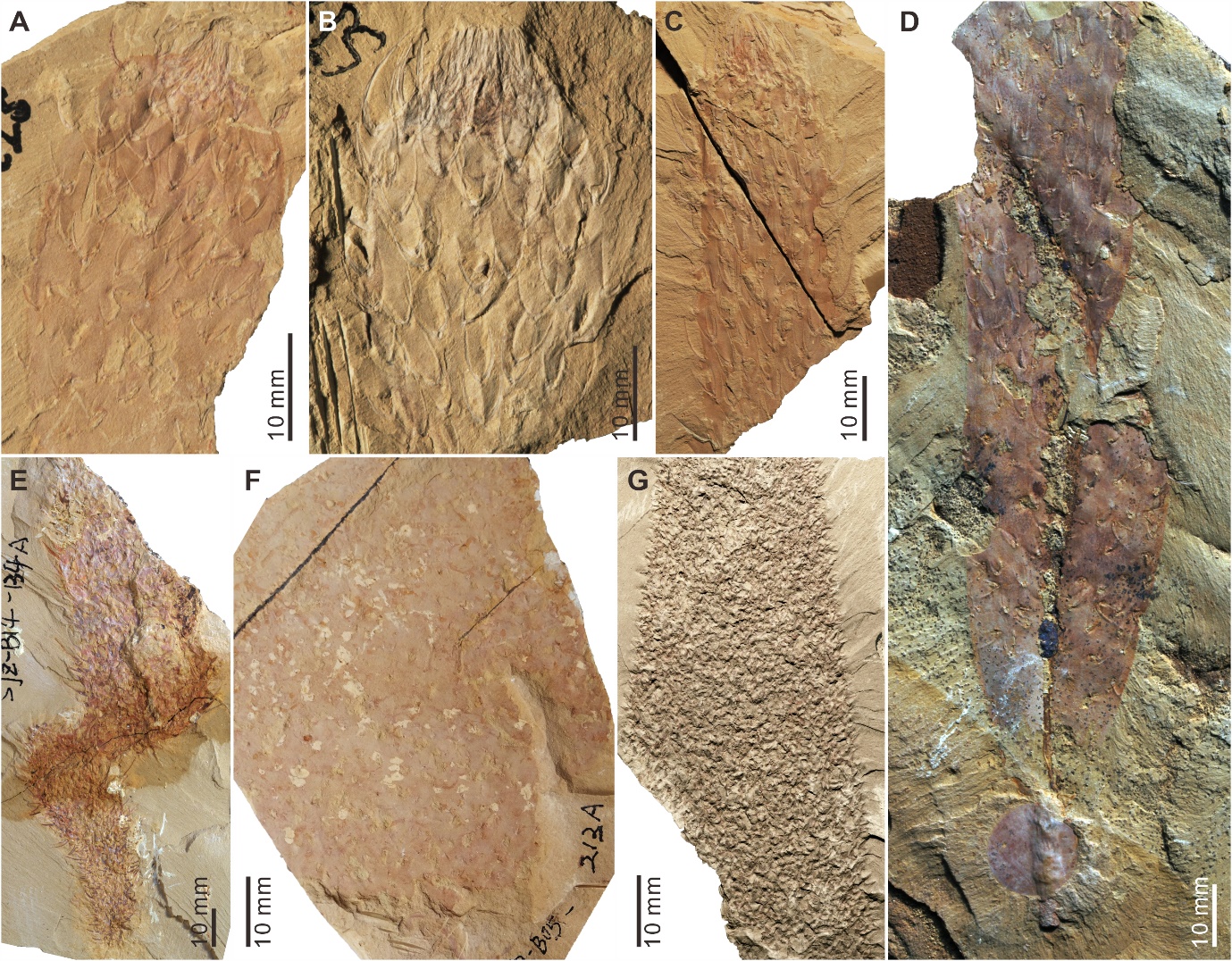

**FIG. S1.** Representative chancelloriids from the Cambrian Chengjiang biota in Kunming area, Yunnan Province, China. (**A–D**) *Allonnia phrixothrix* Bengtson and Hou, 2001; (**A**) SJZ-B04-095B; (**B**) SJZ-B07-115B (modified after Yun et al., 2018, Fig. 4A); (**C**) SJZ-B02-176A; (**D**) SJZ-B03-362, showing a distinctive basal stalk. (**E**) *Allonnia erjiensis* Yun, Zhang, and Li, 2018; SJZ-B14-134 (modified after Yun et al., 2018, Fig. 4C). (**F**) *Allonnia nuda* Cong et al., 2018; SJZ-B05-213A. (**G**) *Dimidia simplex* Jiang in Luo et al., 1982; MF-001A (modified after Yun et al., 2024, Fig. 3E).

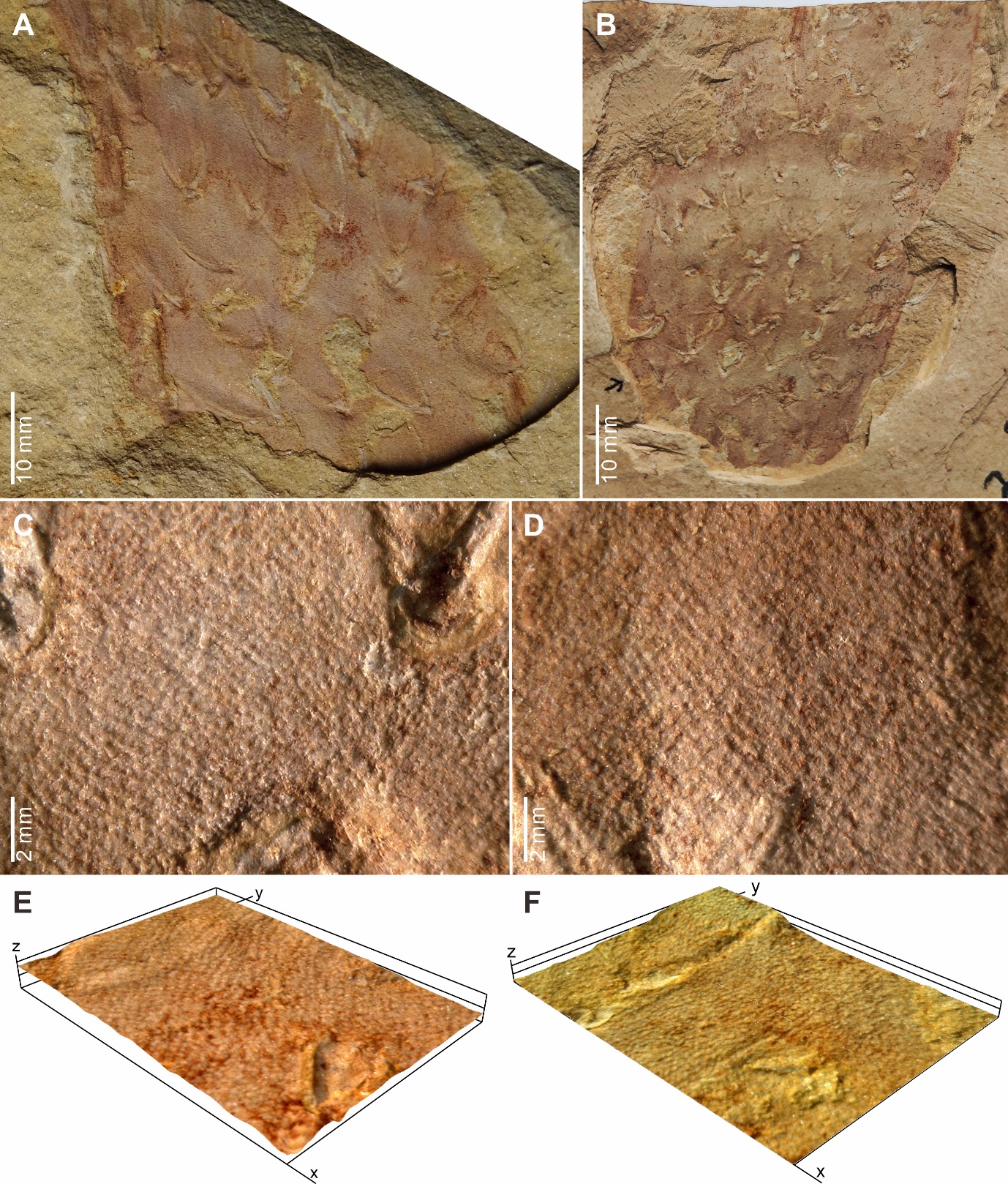

**FIG. S2.** Microstructures of the chancelloriid integument. (**A** and **B**) *Allonnia phrixothrix* from the Chengjiang biota, specimens SJZ-B05-010A and SJZ-B04-461A, respectively. (**C** and **D**) Details of (**B**), showing remarkable protuberances, some of which are arranged in lines to form couples of stripes and grooves. (**E** and **F**) Composite 3-dimensional digital photos of the soft integument, showing protuberances (**E**) and wrinkles (**F**) that corresponding to Figures 1G and 2D, respectively.

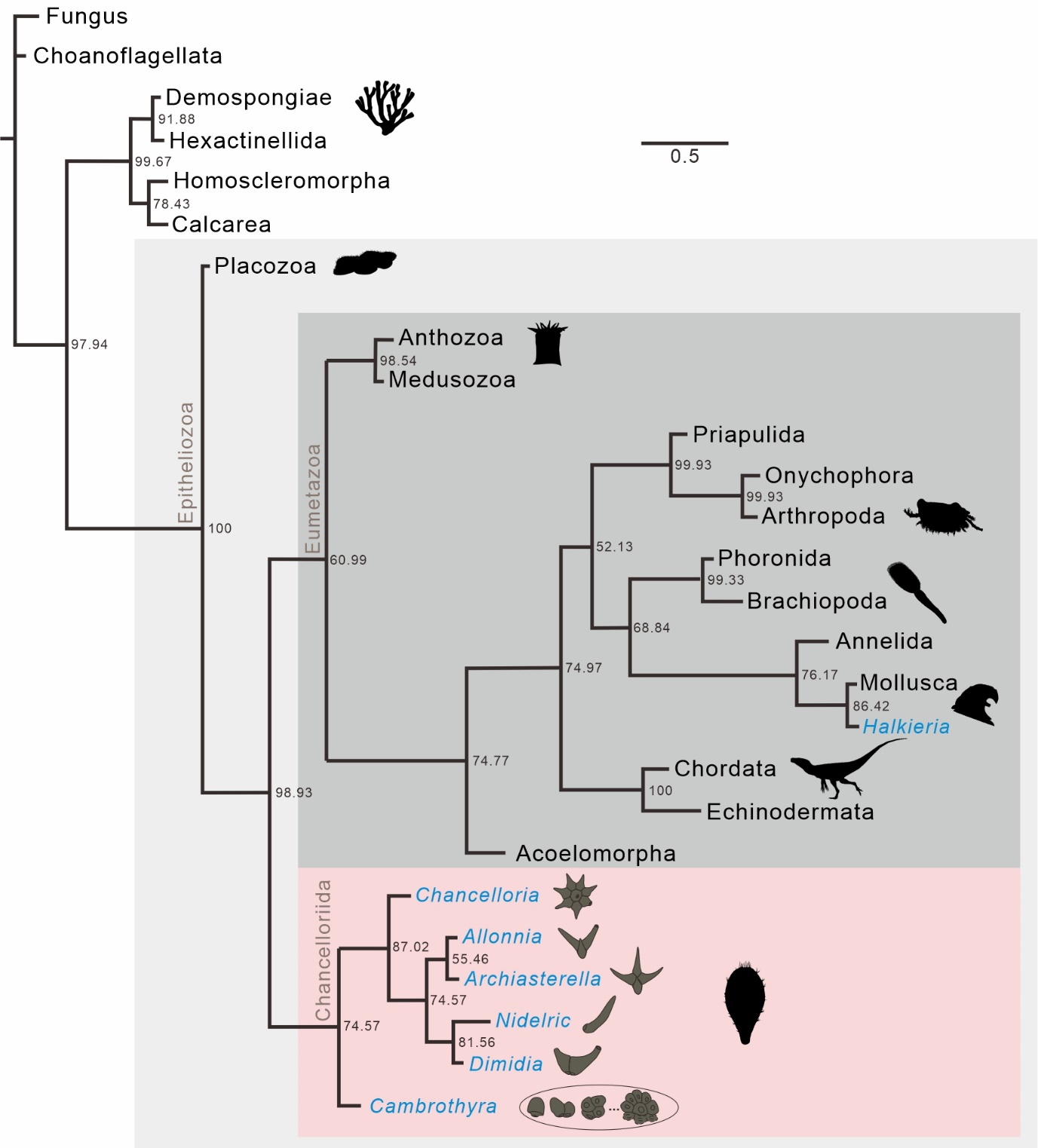

**FIG. S3.** The original phylogenetic tree obtained in the Bayesian cladistic analysis (see Data S2 for character matrix). The analysis was performed by using maximum parsimony with equal character weights and the maximum likelihood and gamma-distributed rate (mki + Γ) model in MrBayes 3.2.7. The consensus tree was resolved from credible sets of 659 sampled trees, based on a dataset containing 26 taxa and 117 characters. Numbers at the nodes are percentages of posterior probabilities, and the scale bar indicates the unit of the expected number of substitutions per site. Silhouettes of the representative animal clades are from PhyloPic (www.phylopic.org), except the one for chancelloriids. Within the clade of chancelloriids, the grey sketches are typical sclerite forms of each genus.

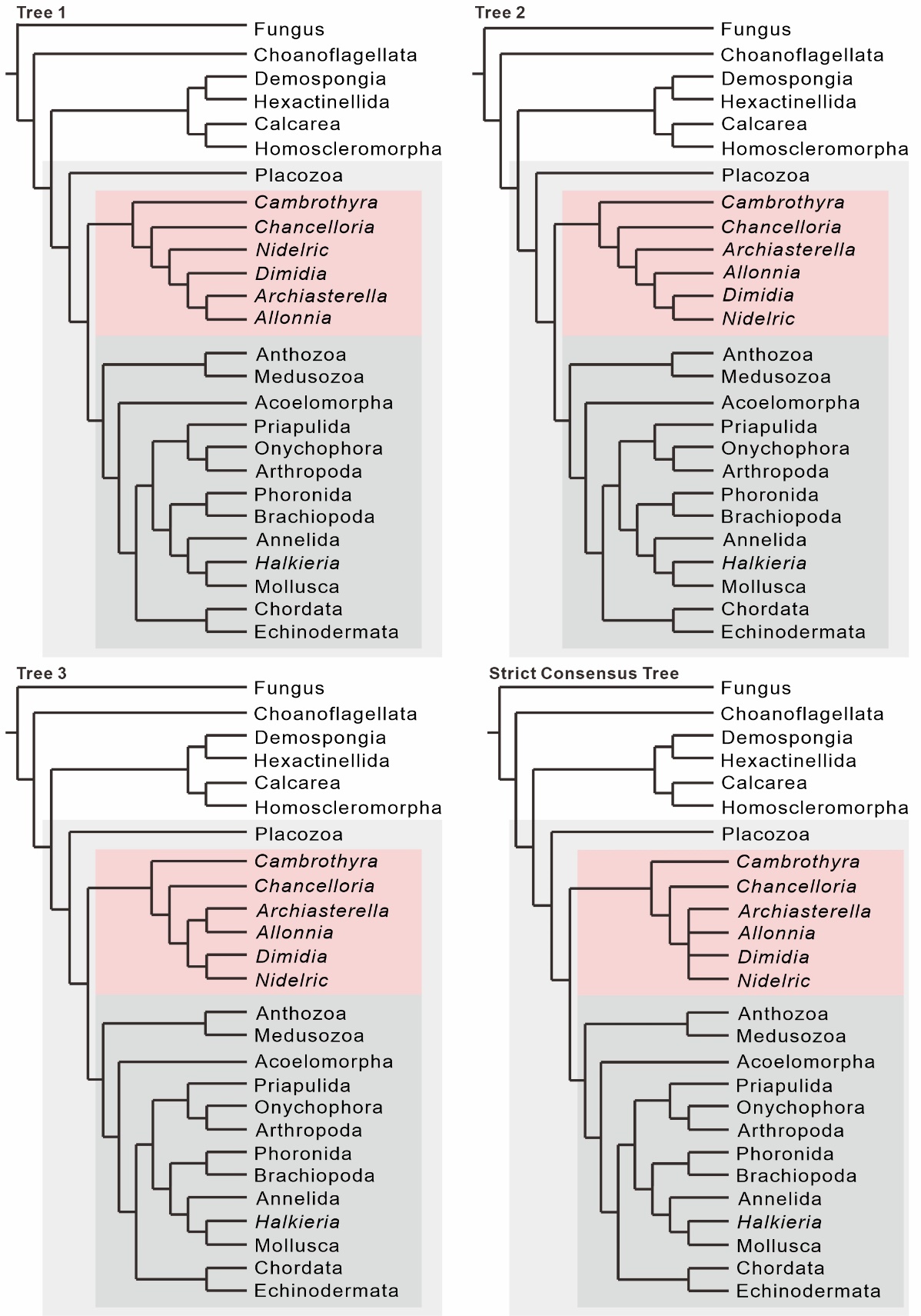

**FIG. S4.** Phylogenetic trees obtained in the TNT cladistic analysis (see Data S2 for character matrix), including 3 maximum parsimonious trees 1 to 3 (tree length = 148; total fit = 93.8; adjusted homoplasy = 7.2, for all the trees) and a strict consensus tree of them.

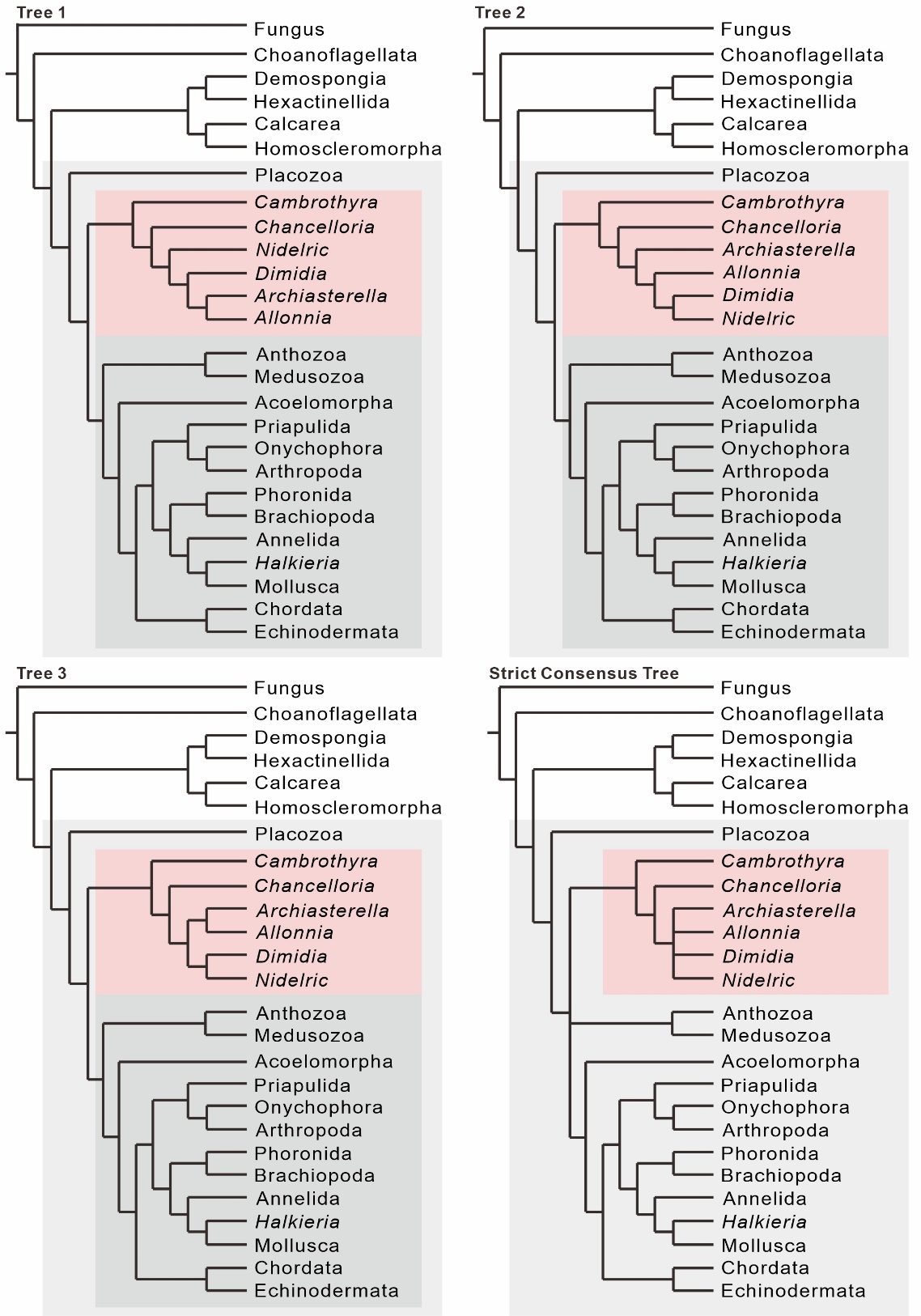

**FIG. S5.** Phylogenetic trees obtained in the additional cladistic analysis based on a slightly modified character matrix (Data S3), a ‘conservative’ version with more “?” coding for chancelloriids (see Appendix S2 for detail), from the one in the main cladistic analysis of this study. The results include 3 maximum parsimonious trees 1 to 3 (tree length = 148; total fit = 93.8; adjusted homoplasy = 7.2) and a strict consensus tree of them.
