## Appendix; Data for "An extinct clade sister to Eumetazoa: On the phylogeny of the Cambrian chancelloriids": Supp.file.2_Appendix.docx

**Appendix S1. Description on the character matrix for the cladistic analyses**

In all characters: ‘0’ = absent, ‘1’ = present, ‘-’ = not applicable, unless otherwise stated. Program files for cladistic analysis are provided as a separated file (Data S2).

1) Multicellularity (Peterson & Eernisse 2001). ‘1’ for chancelloriids.
2) Body symmetry (Philippe *et al.* 2009; Borchiellini *et al.* 2021). ‘1’ for chancelloriids. Chancelloriids have a cylindrical, radially symmetrical body (Bengtson & Collins 2015; Yun *et al.* 2018; Yun & Brock *et al.* 2019; Peng *et al.* 2024).
3) Body tissue (or body regionalization) (Dunn *et al.* 2021). ‘1’ for chancelloriids, since they at least have tissue-or organ-grade body part for biomineralization.
4) Choanocytes (Peterson & Eernisse 2001). ‘0’ for chancelloriids.
5) Pinacocytes (Philippe *et al.* 2009). ‘0’ for chancelloriids.
6) Collar cell complex (Brunet & King 2017). ‘?’ for chancelloriids.
7) Siliceous skeletons (spicules) (Peterson & Eernisse 2001; Philippe *et al.* 2009; Borchiellini *et al.* 2021). ‘0’ for chancelloriids.
8) Calcareous skeletons (Murdock 2020). ‘1’ for chancelloriids and *Halkieria*. Sclerites of chancelloriids and *Halkieria* were originally composed of fibrous aragonites (Bengtson 2005; Porter 2008; Yun & Zhang *et al.* 2021).
9) Poriferan aquiferous system (ostia + oscula) (Peterson & Eernisse 2001; Philippe *et al.* 2009). ‘0’ for chancelloriids except for *Cambrothyra* (‘?’ for this isolated sclerites-based taxon since there are no complete bodies ever found).
10) Sessile filter feeding (adults). ‘1’ for chancelloriids except for *Cambrothyra* (‘?’ for this isolated sclerites-based taxon since there are no complete bodies ever found).
11) Basement membrane (Peterson & Eernisse 2001; Philippe *et al.* 2009). ‘1’ for chancelloriids. Chancelloriids have basement membrane because they have well-developed epidermis.
12) Specialized and reinforced epidermis. ‘1’ for chancelloriids. The strengthening and flexible integument (epidermis) of chancelloriids are functioned in supporting the body and probably related to a cuticular or cuticle-like composition; part of the epidermis are specialized to serve in the biomineralization (Bengtson & Hou 2001; Bengtson & Collins 2015; Yun & Zhang *et al.* 2021). This feature is radically different from that of sponges, including the fragile ‘epidermis’ of homoscleromorphs.
13) Gland cells in epithelia (Martindale 2005; Philippe *et al.* 2009; DuBuc *et al.* 2019). ‘1’ for chancelloriids, refering to the restored biomineralization model for chancelloriids (Bengtson & Hou 2001; Porter 2008; Yun & Zhang *et al.* 2021).
14) Neuro-sensory system (Peterson & Eernisse 2001; Philippe *et al.* 2009). ‘0’ for chancelloriids except for *Cambrothyra* (‘?’ for this isolated sclerites-based taxon).
15) Circular and longitudinal muscles (Peterson & Eernisse 2001). ‘0’ for chancelloriids except for *Cambrothyra* (‘?’ for this isolated sclerites-based taxon). The absence of tentacles or other locomotive organs indicates that chancelloriids don’t have a developed muscular system.
16) Epithelio-muscle cells (EMCs) (Schmidt-Rhaesa 2007). ‘1’ for chancelloriids except for *Cambrothyra* (‘?’ for this isolated sclerites-based taxon), based on the protuberances and stripes (wrinkles) on the soft body wall (see main text).
17) Mesoderm (Peterson & Eernisse 2001; Martindale 2005; Philippe *et al.* 2009). ‘?’ for chancelloriids.
18) Bilaterality (of adults) (Peterson & Eernisse 2001; Philippe *et al.* 2009). ‘0’ for chancelloriids except for *Cambrothyra* (‘?’ for this isolated sclerites-based taxon).
19) Anterior/posterior axis (Martindale 2005). ‘?’ for chancelloriids. The anterior/posterior axis is present in the larva of sponges (Wörheide *et al.* 2012). The adults of chancelloriids don’t have an anterior/posterior axis like sponges, but there is no knowledge about their larva or early development.
20) Oral/aboral axis (Martindale 2005; DuBuc *et al.* 2019). ‘1’ for chancelloriids except for *Cambrothyra* (‘?’ for this isolated sclerites-based taxon). The conspicuous apical orifice indicates an oral orientation.
21) Dorsal/ventral axis (Peterson & Eernisse 2001; Martindale 2005; DuBuc *et al.* 2019). ‘?’ for chancelloriids. The adults of chancelloriids don’t have a dorsal/ventral axis, but there is no knowledge about their larva or early development.
22) Dorsal/ventral polarity (Martindale 2005; DuBuc *et al.* 2019). ‘?’ for chancelloriids. The adults of chancelloriids don’t have a dorsal/ventral polarity, but there is no knowledge about their larva or early development.
23) Segmentation of ectodermal and mesodermal structures (Peterson & Eernisse 2001; Blair 2008). ‘0’ for chancelloriids except for *Cambrothyra* (‘?’ for this isolated sclerites-based taxon).
24) Through-gut (Martindale 2005). ‘0’ for chancelloriids except for *Cambrothyra* (‘?’ for this isolated sclerites-based taxon). There is no internal gut in the chancelloriid body.
25) Tentacles (Han *et al.* 2016). ‘0’ for chancelloriids except for *Cambrothyra* (‘?’ for this isolated sclerites-based taxon).
26) Cnidocil (Han *et al.* 2016). ‘0’ for chancelloriids except for *Cambrothyra* (‘?’ for this isolated sclerites-based taxon) since there are no tentacles.
27) Lophophore (Peterson & Eernisse 2001). ‘0’ for chancelloriids except for *Cambrothyra* (‘?’ for this isolated sclerites-based taxon).
28) Ciliated extensions of the mesocoel (Peterson & Eernisse 2001). ‘0’ for chancelloriids except for *Cambrothyra* (‘?’ for this isolated sclerites-based taxon) and ‘?’ for *Halkieria*.
29) Prototroch (Peterson & Eernisse 2001). Larval character. ‘?’ for chancelloriids and *Halkieria*.
30) Metatroch (Peterson & Eernisse 2001). Larval character. ‘?’ for chancelloriids and *Halkieria*.
31) Adoral ciliary band (Peterson & Eernisse 2001). Larval character. ‘?’ for chancelloriids and *Halkieria*.
32) Telotroch (Peterson & Eernisse 2001). Larval character. ‘?’ for chancelloriids and *Halkieria*.
33) Trimery (Peterson & Eernisse 2001). Larval character. ‘?’ for chancelloriids and *Halkieria*.
34) Metanephridia open through metacoel (Peterson & Eernisse 2001). Larval character. ‘?’ for chancelloriids and *Halkieria*.
35) Metanephridia with coelomic compartment restricted to sacculus (Peterson & Eernisse 2001). Larval character. ‘?’ for chancelloriids and *Halkieria*.
36) Hydropore (Peterson & Eernisse 2001). Larval character. ‘?’ for chancelloriids and *Halkieria*.
37) Inner epithelium secreting periostracum (Peterson & Eernisse 2001). ‘?’ for chancelloriids and *Halkieria*.
38) Trilayered cuticle (Peterson & Eernisse 2001). ‘?’ for chancelloriids and *Halkieria*.
39) Ecdysis (Peterson & Eernisse 2001). ‘0’ for chancelloriids except for *Cambrothyra* (‘?’ for this isolated sclerites-based taxon).
40) Setae (Peterson & Eernisse 2001). ‘0’ for chancelloriids and ‘1’ for *Halkieria* (Vinther *et al.* 2017).
41) Protrusible and retractible setae (Peterson & Eernisse 2001). ‘-’ for chancelloriids except for *Cambrothyra* (‘?’ for this isolated sclerites-based taxon) and ‘?’ for *Halkieria* (Vinther *et al.* 2017).
42) Radula. ‘0’ for chancelloriids except for *Cambrothyra* (‘?’ for this isolated sclerites-based taxon) and ‘1’ for *Halkieria* (representing the ‘halwaxiid’ group) (Vinther *et al.* 2017).
43) Head divided into three segments (Peterson & Eernisse 2001). ‘-’ for chancelloriids except for *Cambrothyra* (‘?’ for this isolated sclerites-based taxon).
44) Terminal mouth (Peterson & Eernisse 2001). ‘0’ for chancelloriids except for *Cambrothyra* (‘?’ for this isolated sclerites-based taxon).
45) Food modified with limbs (Peterson & Eernisse 2001). ‘-’ for chancelloriids except for *Cambrothyra* (‘?’ for this isolated sclerites-based taxon).
46) Anus (Peterson & Eernisse 2001). ‘0’ for chancelloriids except for *Cambrothyra* (‘?’ for this isolated sclerites-based taxon).
47) Pharyngotremy (Peterson & Eernisse 2001). ‘0’ for chancelloriids except for *Cambrothyra* (‘?’ for this isolated sclerites-based taxon).
48) Pharyngeal gill slits (Peterson & Eernisse 2001). ‘-’ for chancelloriids except for *Cambrothyra* (‘?’ for this isolated sclerites-based taxon).
49) Ventral nervous system (Peterson & Eernisse 2001). ‘0’ for chancelloriids except for *Cambrothyra* (‘?’ for this isolated sclerites-based taxon).
50) Circumoesophageal nerve ring (Peterson & Eernisse 2001). ‘0’ for chancelloriids except for *Cambrothyra* (‘?’ for this isolated sclerites-based taxon).
51) Diffuse nerve system (Peterson & Eernisse 2001; Han *et al.* 2016). ‘?’ for chancelloriids. It is impractical to prove whether chancelloriids have diffuse nerve system or not.
52) Glial interstitial cell system (Peterson & Eernisse 2001). ‘?’ for chancelloriids.
53) Nerve cells organized into distinct ganglia (Peterson & Eernisse 2001). ‘0’ for chancelloriids except for *Cambrothyra* (‘?’ for this isolated sclerites-based taxon). There is no trace of a ganglia even in the best-preserved chancelloriid bodies.
54) Endoskeleton mesodermally derived (Peterson & Eernisse 2001). ‘0’ for chancelloriids.

*Cellular, genetic, and developmental level characters that frame the animal phylogenetic tree (Peterson & Eernisse 2001) (‘?’ for chancelloriids and mostly ‘?’ for Halkieria):*

55) Septate junctions.
56) Gap junctions.
57) Hemidesmosomes.
58) Flagellar vanes.
59) Incubated cinctoblastula larva with cross-striated ciliary rootlets (Philippe *et al.* 2009).
60) ‘Acoelomorph’ type of ciliary rootlet.
61) Belt desmosomes.
62) Basal lamina.
63) Ciliated epidermis.
64) Densely multiciliated epidermis.
65) Distinct ‘step’ in cilia.
66) Egg with four polar bodies.
67) Acrosome (Ax 1996). 0 absent; 1 present; 2 present as a distinct organelle.
68) Perforatorium (subacrosomal material).
69) Germ line and gonads.
70) Gonads present with gametes passing through coelom and metanephridium.
71) One axis prespecified during oogenesis.
72) Stereotypical cleavage pattern.
73) Spiral cleavage with 4d mesoderm.
74) Gastrulation.
75) Blastopore associated with larval/adult mouth.
76) Blastopore associated with larval/adult anus.
77) Lophotrochozoan 18S rDNA.
78) Endomesodermal muscle cells.
79) Endomesoderm derived from gut.
80) Ectomesenchyme.
81) 4d endomesoderm.
82) Mesodermal germ bands derived from 4d.
83) Lateral coelom derived from mesodermal bands.
84) Somatoblast.
85) Podocytes/terminal cells/nephrocytes.
86) Cuticle with chitin.
87) Nerve cells.
88) Acetylcholine used as a neurotransmitter.
89) Brachyury expressed in oral and anal regions of gut.
90) Hemerythrin.
91) Nuclear lamins.
92) Intermediate filament proteins: 0 = absent; 1 = present as S-type (deletion of 42 residues and laminin similarity region absent); 2 = present as L-type (presence of 42 residues in coil 1b subdomain and laminin similarity tail).
93) tRNA Lys.
94) AUA methionine.
95) AGA and AGG serine.
96) Hox/Parahox genes (Peterson & Eernisse 2001; Jakob *et al.* 2004; Martindale 2005).
97) Hox complex consisting of seven genes.
98) Central Hox class member(s).
99) Antp.
100) Ubx/abd-A.
101) Lox2/4.
102) Hox 6–8.
103) Abd-B duplication (Lophotrochozoa).
104) Abd-B duplication (Deuterostomia).
105) Hexapeptide.
106) T-box genes.

*Characters of chancelloriids (Luo et al. 1982; Beresi & Rigby 1994; Bengtson & Hou 2001; Janussen et al. 2002; Beresi 2003; Bengtson 2005; Randell et al. 2005; Porter 2008; Kloss et al. 2009; Beresi & Rigby 2013; Hou et al. 2014; Bengtson & Collins 2015; Cong et al. 2018; Yun et al. 2018; Yun & Brock et al. 2019; Yun & Zhang et al. 2019; Yun & Cui et al. 2021; Yun & Zhang et al. 2021; Yun et al. 2024), Halkieria (Bengtson & Missarzhevsky 1981; Conway Morris & Peel 1995; Conway Morris & Chapman 1996; Qian et al. 1999; Porter 2004; Bengtson 2005; Conway Morris & Caron 2007; Porter 2008), and some mollusks (Vinther et al. 2017):*

107) Aesthete canal system. ‘0’ for chancelloriids except for *Cambrothyra* (‘?’ for this isolated sclerites-based taxon), ‘1’ for *Halkieria* and Mollusca.
108) Coelosclerites (hollow mineralized sclerites). ‘1’ for chancelloriids and *Halkieria*, ‘0/1’ for Mollusca.
109) Sclerites composed of hollow long conical rays. ‘1’ for common chancelloriids, ‘0’ for *Cambrothyra* and *Halkieria*.
110) Sclerites composed of hollow short conical (elliptical or drop-shaped) rays. ‘1’ for chancelloriids and ‘0’ for *Halkieria*.
111) Coelosclerites densely covered the body. ‘1’ for *Nidelric*, *Dimidia*, and *Halkieria*; ‘0/1’ for *Allonnia* (at one species *Al. erjiensis* has dense sclerites); ‘?’ for *Cambrothyra*.
112) Uniform coelosclerites (except for the modified sclerites in apical tuft) in a scleritome. ‘1’ for only *Allonnia*, *Archiasterella*, *Nidelric*, and *Dimidia*; ‘0’ for Chancelloria, *Cambrothyra*, and *Halkieria*.
113) Composite coelosclerites that composed of two or more hollow rays. ‘1’ for chancelloriids except for *Nidelric*.
114) Coelosclerites have lateral rays. ‘1’ for all chancelloriids except for *Nidelric*.
115) Coelosclerites have central rays. ‘1’ for *Chancelloria* and *Cambrothyra*; ‘0’ for other chancelloriids.
116) Coelosclerites have ascending rays. ‘1’ for only *Allonnia*, *Archiasterella*, and *Cambrothyra*; ‘0’ for other chancelloriids.
117) Apical tuft composed of modified single-rayed sclerites. ‘1’ for all chancelloriids except for *Cambrothyra* (‘?’ for this isolated sclerites-based taxon).

**Appendix S2. Character modifications in the matrix for the additional cladistic analysis**

The additional analysis is based on a slightly modified character matrix (see Data S3, which is a ‘conservative’ version with more “?” coding for chancelloriids) from the one in the main cladistic analysis of this study. In the character matrix:

Character 4 (Choanocytes): “?” for chancelloriids.

Character 5 (Pinacocytes): “?” for chancelloriids.

Character 9 (Aquiferous system): “?” for chancelloriids.

Character 14 (Neuro-sensory system): “?” for chancelloriids.

Character 16 (EMCs): “?” for chancelloriids.

Character 26 (Cnidocil): “?” for chancelloriids.

The analysis was operated in TNT 1.5 (with a ‘traditional search’ algorithm), resolving three maximum parsimonious trees (tree length = 148; total fit = 93.8; adjusted homoplasy = 7.2) and a strict consensus tree of them (Fig. S5).
